# Auditory Brainstem Responses in the C57BL/6J Fragile X Syndrome Knockout Mouse Model

**DOI:** 10.1101/2021.10.27.466191

**Authors:** Amita Chawla, Elizabeth A McCullagh

## Abstract

Sensory hypersensitivity, especially in the auditory system, is a common symptom in Fragile X Syndrome (FXS), the most common monogenic form of intellectual disability. However, linking phenotypes across genetic background strains of mouse models has been a challenge and could underly some of the issues with translatability of drug studies to the human condition. This study is the first to characterize the auditory brainstem response (ABR), a minimally invasive physiological readout of early auditory processing that is also used in humans, in a commonly used mouse background strain model of FXS, C57BL/6J. We measured morphological features of pinna and head and used ABR to measure hearing range, monaural and binaural auditory responses in hemizygous males, homozygous females and heterozygous females compared to wildtype mice. Consistent with previous work we showed no difference in morphological parameters across genotypes or sexes. Male FXS mice had increased threshold for high frequency hearing at 64 kHz compared to wildtype males, while females had no difference in hearing range between genotypes. In contrast, female homozygous FXS mice had decreased amplitude of wave IV of the monaural ABR, while there was no difference in males for amplitudes and no change in latency of ABR waveforms across sexes and genotypes. Lastly, FXS males had increased latency of the binaural interaction component (BIC) at 0 ITD compared to wildtype males. These findings further clarify auditory brainstem processing in FXS by adding more information across genetic background strains allowing for a better understanding of shared phenotypes.

## 1 Introduction

Fragile X Syndrome (FXS) is the most common monogenic form of autism spectrum disorder (ASD) and shares many attributes of ASDs including auditory hypersensitivity and other sensory disruptions (Abbeduto and Hagerman, 1997; Chen and Toth, 2001; Hagerman and Hagerman, 2002; Arnett et al., 2014). FXS is a tractable genetic model for ASD with several commercially available models, including rat and mouse (The Dutch-Belgian Fragile X Consorthium et al., 1994; Till et al., 2015; Tian et al., 2017). Despite the common use of these models to study FXS, phenotypes are not always shared between species and background strains, particularly for sensory processing. As a result, drug therapies have struggled to rescue the human disorder (Dahlhaus, 2018). One of the most common symptoms described in people with FXS and ASD is auditory hypersensitivity (Ethridge et al., 2017; Stefanelli et al., 2020). The mechanisms that underly auditory alterations are unknown, but likely involve the entirety of the ascending pathway from the periphery to the cortex (reviewed in McCullagh et al., 2020b). A complete characterization of auditory processing from the periphery to cortex across sexes, background strains, and models is needed to fully understand shared phenotypes and circuitry involved in this common symptom.

The auditory brainstem is one brain region in the ascending auditory pathway that has shown to have anatomical, physiological, and behavioral alterations in FXS mouse models (Brown et al., 2010; Beebe et al., 2014; Wang et al., 2014, 2015; Rotschafer et al., 2015; Garcia-Pino et al., 2017; McCullagh et al., 2017, 2020a; Rotschafer and Cramer, 2017; Curry et al., 2018; El-Hassar et al., 2019; Lu, 2019). The auditory brainstem is the first site of binaural processing of sound location in the brain using interaural timing and level differences (ITD and ILD respectively) to compute sound source locations (Grothe et al., 2010). This brain area is also involved in separating spatial channels allowing for complex listening environments. Disruptions in this spatial separation and binaural processing could lead to auditory hypersensitivity due to inability to separate sound sources (Bronkhorst, 2015). One measure of auditory brainstem physiology, and binaural hearing, that can be directly translated between animal models and humans is the auditory brainstem response (ABR) (Laumen et al., 2016).

The ABR is a minimally invasive physiological measure that allows for simultaneous assessment of sound processing across multiple brainstem nuclei, as each wave of the ABR directly corresponds to distinct areas of the ascending auditory brainstem pathway. These features make the ABR an attractive translational tool. Indeed, recent evidence suggests that ABR measurements are an early indicator for auditory dysfunction in ASD (Santos et al., 2017). ABRs can also be used to assess binaural hearing, which is essential for sound localization and hearing in noisy environments and often impaired in ASD (Visser et al., 2013). Monoaural ABRs can be recorded by stimulating each ear separately and binaural responses can be generated by stimulating both ears simultaneously. The sum of the two monaural (left and right) responses should equal the binaural (both ear) response since the recruited neural activity from each ear should be double when stimulated simultaneously. However, this is not the case, there is a difference that arises when the summed monoaural responses are subtracted from the binaural response, called the binaural interaction component (BIC). The BIC is thought to be a direct measure of binaural processing ability in humans and animals that requires the precise balance of excitatory and inhibitory drive in brainstem sound localization circuits (Laumen et al., 2016).

In this study we report on the hearing ability, using the ABR and morphological craniofacial and pinna features, of the most common mouse model of FXS, C57BL/6J across the sexes and females heterozygous for the Fmr1 mutation. We hypothesize that there may be sex differences in ABRs independent of FXS genotype, but that additionally FXS animals are likely to have alterations in peak amplitude or latency of ABRs and impaired high frequency hearing compared to wildtype consistent with work in other FXS mouse strains (Kim et al., 2013; Rotschafer et al., 2015; El-Hassar et al., 2019). Establishing core auditory phenotypes across the sexes and different mouse strains is key to creating a toolbox of techniques that may translate to human FXS.

## 2 Materials and Methods

All experiments complied with all applicable laws, National Institutes of Health guidelines, and were approved by the Oklahoma State University IACUC.

### 2.1 Animals

Experiments were conducted in C57BL/6J (stock #000664, B6) wildtype background, hemizygous male, homozygous, or heterozygous female *Fmr1* mutant mice (B6.129P2-*Fmr1*^*tm1Cgr*^/J stock #003025, Fmr1 or Fmr1 het respectively) obtained from the Jackson Laboratory (Bar Harbor, ME USA)(The Dutch-Belgian Fragile X Consorthium et al., 1994). Sex was treated as a biological variable and differences between the sexes, when present, are noted in the results. Numbers of animals for each experiment used are listed in the figure legends and ranged from 6-10 animals per sex and genotype. Animals ranged in age from 62 – 120 days old (average ages per genotype 89 ± 4 days old B6, 101 ± 3 days old Fmr1, and 97 ± 4 days old Fmr1 het).

### 2.2 Morphological Measures

Features of animal’s head, pinna, and body mass (weight) were measured for each genotype using 6 Inch Stainless Steel Electronic Vernier Calipers (DIGI-Science Accumatic digital caliper Gyros Precision Tools Monsey, NY, USA) and an electronic scale. The distance between the two pinnae (inter pinna distance), distance from the nose to the middle of the pinna (nose to pinna distance) and pinna width and length were measured (Figure 2A). The effective diameter was calculated as the square root of pinna length times pinna width (Anbuhl et al., 2017).

### 2.3 ABRs

ABR recordings were performed using similar methods to previously published work (Benichoux et al., 2018; McCullagh et al., 2020a; New et al., 2021). Animals were anesthetized using two mixtures of ketamine-xylazine 60mg/kg ketamine and 10mg/kg xylazine for initial induction followed by maintenance doses of 25 mg/kg ketamine and 12 mg/kg xylazine. Once anesthesia was confirmed by lack of a toe-pinch reflex, animals were transferred to a small sound attenuating chamber (Noise Barriers Lake Forest, IL, USA) and body temperature was maintained using a water-pump heating pad. Subdermal needle electrodes were placed under the skin between the ears (apex), directly behind the apex in the nape (reference), and in the back leg for ground. This montage has been shown to be particularly effective in generating the BIC (Levine, 1981; Laumen et al., 2016). Evoked potentials from subdermal needle electrodes were acquired and amplified using a Tucker-Davis Technologies (TDT, Alachua, FL, USA) RA4LI head stage and a TDT RA16PA preamplifier. Further amplification was provided by a TDT Multi I/O processor RZ5 connected to a PC with custom Python software for data recording. Data were averaged across 500-1000 repetitions per condition and processed using a second order 50 - 3000 Hz filter over 12 ms of recording time.

Sound stimuli (see below for varying types) were presented to the animal through TDT EC-1 electrostatic speakers (frequencies 32 – 64 kHz) or TDT MF-1 multi-field speakers (frequencies 1 – 24 kHz and clicks) coupled through custom ear pieces fitted with Etymotic ER-7C probe microphones (Etymotic Research Inc, Elk Grove Village, IL, USA) for in-ear calibration (Beutelmann et al., 2015). Sounds were generated using a TDT RP2.1 Real-Time processor controlled by custom Python code at a sampling rate of 48828.125 Hz. Sounds were presented at an interstimulus interval of 30 ms with a standard deviation of 5 ms (Laumen et al., 2016). An additional rejection threshold was set to eliminate high amplitude heart rate responses from average traces and improve signal to noise ratio.

#### 2.3.1 Audiogram

Hearing range of animals was tested using the threshold for hearing across different frequencies of sound (1, 2, 4, 8, 16, 24, 32, 46, 64 kHz). Threshold was determined using a visual detection method (Brittan-Powell and Dooling, 2004), or the lowest level (dB SPL) a response could be detected (independent of wave) or 2.5 dB SPL below the lowest level that elicited a response. Audiogram stimuli consisted of tone bursts (2 ms ± 1 ms on/off ramps) of varying frequency and intensity.

#### 2.3.2 Monaural ABRs

Click stimuli (0.1 ms transient) were presented to each ear independently to generate monaural evoked potentials. Peak amplitude (voltage from peak to trough) and latency (time to peak amplitude) were measured across the four peaks of the ABR waveform at 90 dB SPL (Figure 1A). The trough was considered the lowest point for that wave. Monaural data from the two ears were averaged to determine monaural amplitude and latency for each animal. Like hearing thresholds across frequency, click threshold was determined for each genotype and sex. Click threshold is determined by decreasing intensity of sound in 5 - 10 dB SPL steps until ABR waveforms disappear.

**Figure 1.**
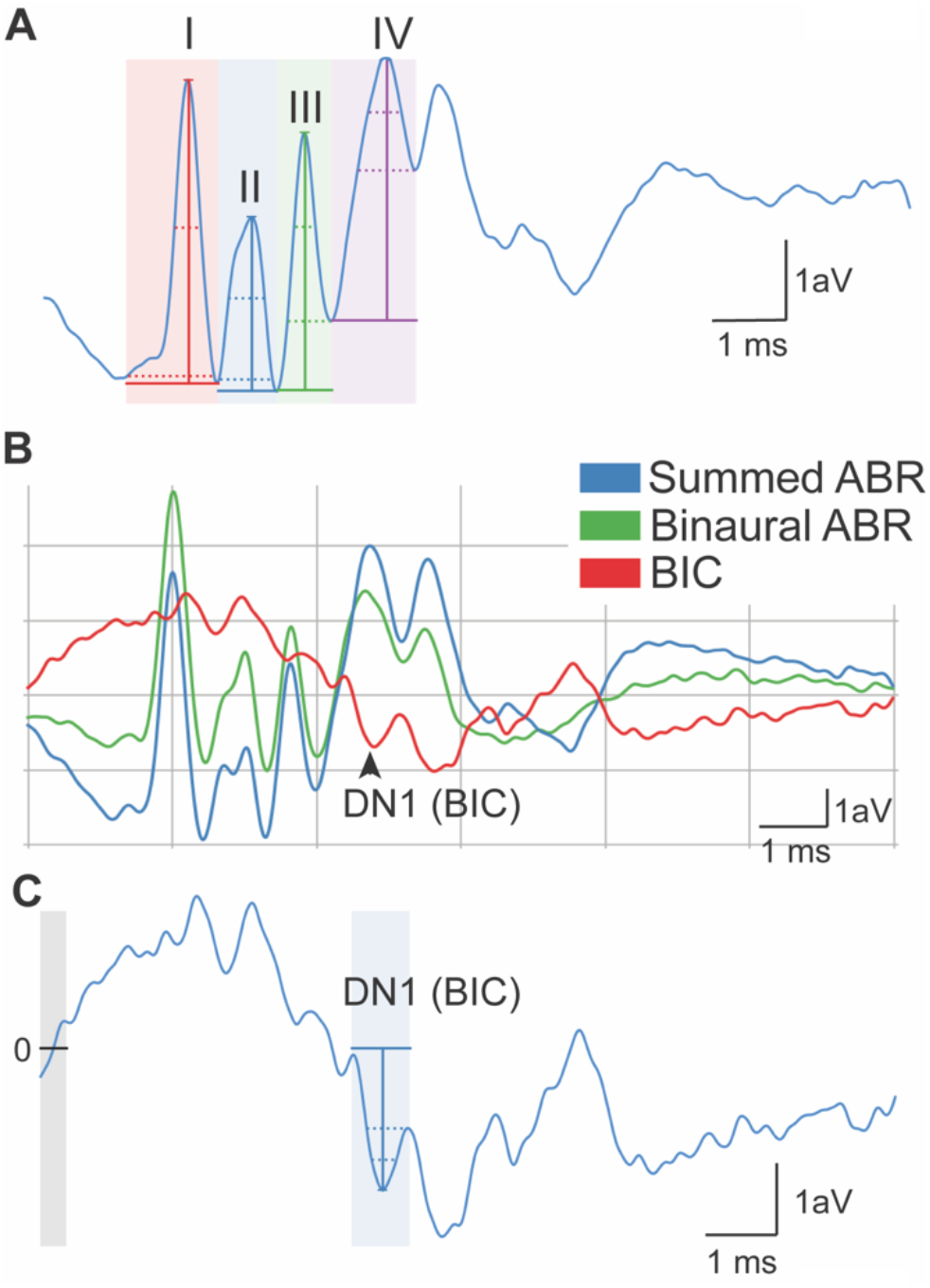
Quantification of ABR signals. Monaural ABR amplitudes were quantified for each ear as the voltage between the peak of the ABR and trough of the waveform for waves I-IV (A). Latency was calculated as the time when the height of the peak occurred. DN1 or BIC (red) was calculated as the prominent negative peak corresponding with wave IV of the summed (blue) and binaural (green)(B). BIC is calculated as the summed ABR subtracted from the binaural ABR. The BIC amplitude was calculated as voltage at the peak of the DN1 waveform to the baseline (0, line and gray area) of the measurement (C). Scale represents 1 voltage unit (Y) during 1 ms (X).

**Figure 2.**
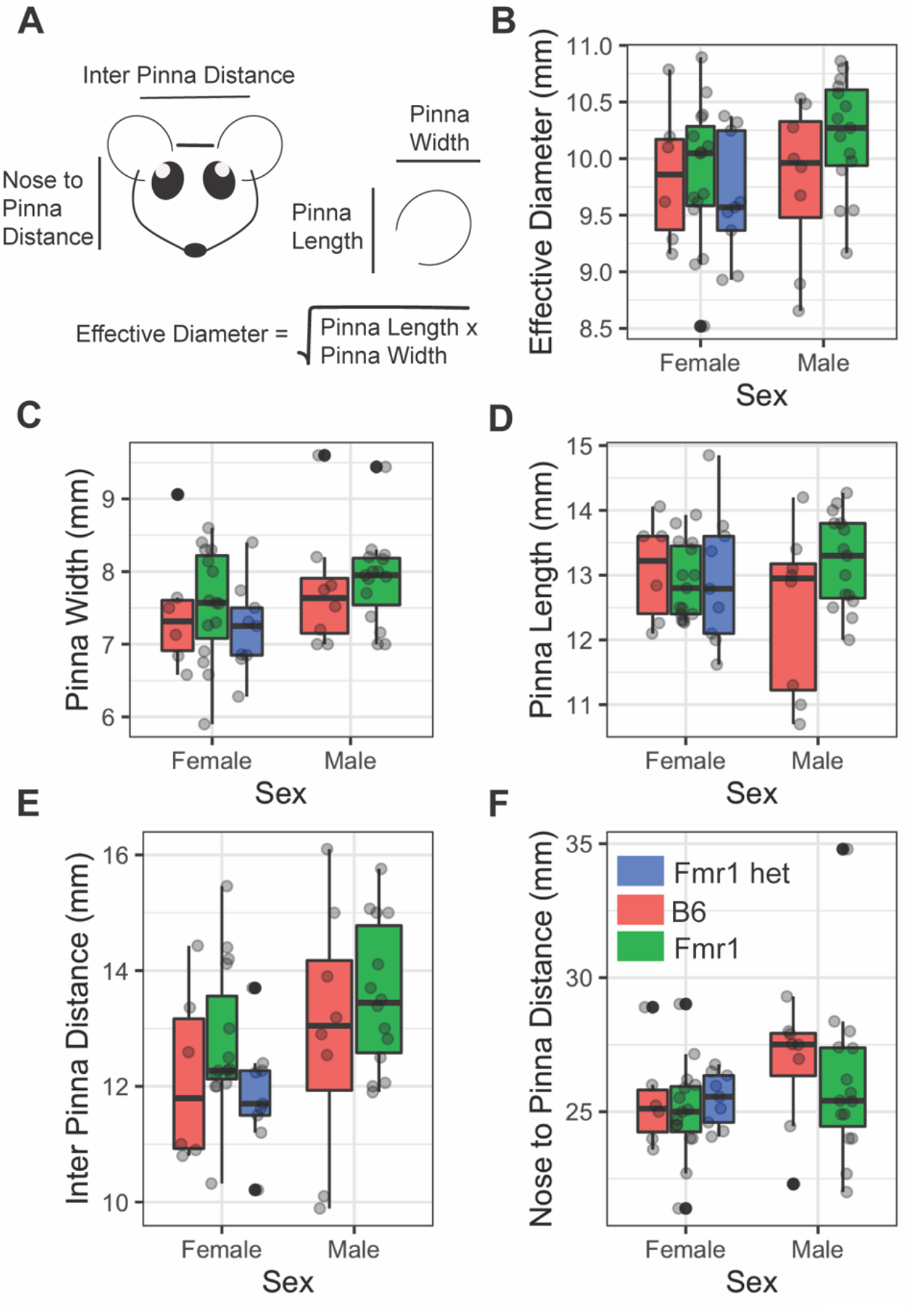
Morphological features of FXS mice. Pinna and head features (A) were measured between the sexes (x-axis) and genotypes (purple = B6, teal = Fmr1, yellow = Fmr1 het). There was no difference between the sexes or genotypes for any of the measures (effective diameter B, pinna width C, pinna length D, inter pinna length E, or nose to pinna length F). Data represent 6 B6, 15 Fmr1, 9 Fmr1 het females and 8 B6, 15 Fmr1 males.

#### 2.3.3 Binaural ABRs

Click stimuli at 90 dB SPL were also presented to both ears simultaneously to generate a binaural evoked potential. The binaural interaction component (BIC) of the ABR was calculated by subtracting the sum of the two monaural ABRs from the binaural ABR (Laumen et al., 2016; Benichoux et al., 2018)(Figure 1B and C). BIC amplitude and latency were then measured using custom Python software, with amplitude being relative to the zero baseline of the measurement (Figure 1C, gray area with line). BIC was characterized as the prominent negative DN1 wave corresponding to the fourth wave of the binaural and summed ABR (Figure 1B). To measure interaural timing difference (ITD) computation using the BIC, animals were presented with stimuli that had varying ITDs of ± 2 ms in 0.5 ms steps and corresponding BIC amplitudes and latencies were calculated like above. This ITD range was chosen to be comparable to other studies in small rodents (Benichoux et al., 2018).

### 2.4 Analysis of ABR waveforms

Custom python software was used to analyze evoked potentials for monaural and binaural stimuli (New et al., 2021). To account for fluctuation in the baseline signal of the ABR, raw traces were zeroed to establish a baseline across traces. The software included automatic peak detection with the capability of manual correction or deselection upon visual confirmation.

### 2.5 Statistical analyses

Figures were generated using R Studio (R Core Team, 2013), ggplot2 (Wickham, 2016), and Adobe Illustrator (Adobe, San Jose, CA USA). Data points on Figures 3, 4, and 5 represent means and error bars reflect standard error, boxplots in Figure 2 display the median and 25^th^ – 75^th^ percentiles (or 1^st^ and 3^rd^ quartiles respectively) the whiskers represent +/-1.5 times the interquartile range. Data that falls outside the range are plotted as individual points. Multivariate data (monaural peak amplitude and latency, audiogram, and BIC amplitude and latency across ITD) were analyzed using linear mixed effects (lme4) models (Bates et al., 2015) with sex, genotype, and condition (ITD, Frequency, Peak) as fixed effects and animal as a random effect. It was expected that there may be differences between the sexes and genotypes therefore a priori, it was determined that estimated marginal means (emmeans; (Lenth, 2019)) would be used for pairwise comparisons between sexes and genotype. Two-way ANOVAs were performed to compare relationships between morphological features, sex, and genotype with adjusted Tukey posthoc analysis to compare groups. Where values are indicated as statistically significant between the two genotypes, * indicated a p-value of < 0.05, ** = p < 0.01, and *** = p < 0.0001.

**Figure 3.**
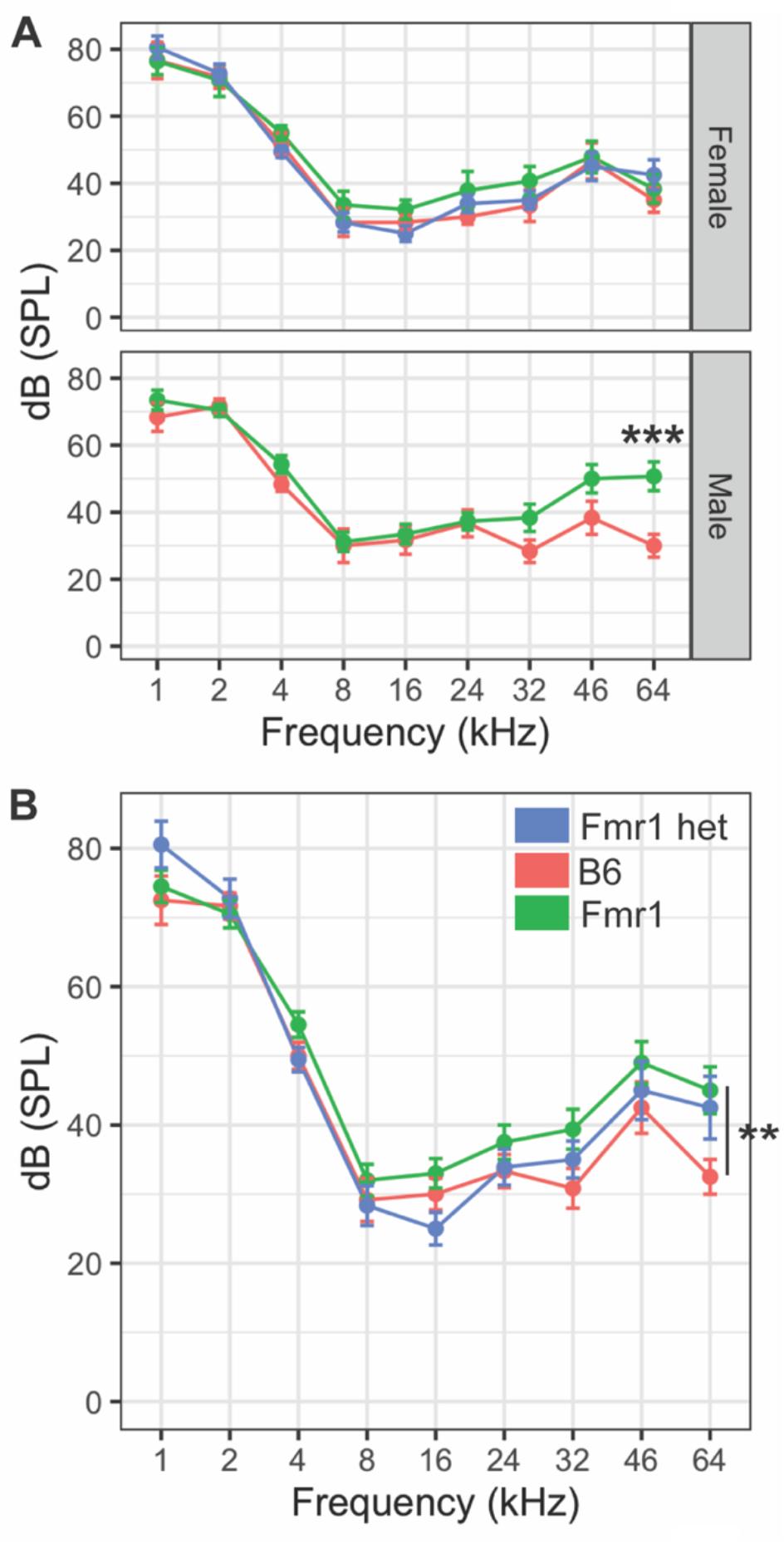
Hearing range of FXS mice. Hearing threshold (dB SPL) was measured across frequencies (1-64 kHz) in male and female mice of all genotypes (A). Fmr1 (green) males (lower panel) had significantly higher threshold hearing at 64 kHz compared to B6 (red) males. There were no differences in hearing range between female Fmr1 (green), B6 (red), and Fmr1 het (blue) mice (top panel A). When sexes were combined, the significant difference in hearing at 64 kHz persisted in Fmr1 animals compared to B6 but was not significant in female Fmr1 het animals (B). ** = p < 0.01, *** = p < 0.001. Data represent 6 B6, 7 Fmr1, 9 Fmr1 het females and 6 B6, 11 Fmr1 males.

**Figure 4.**
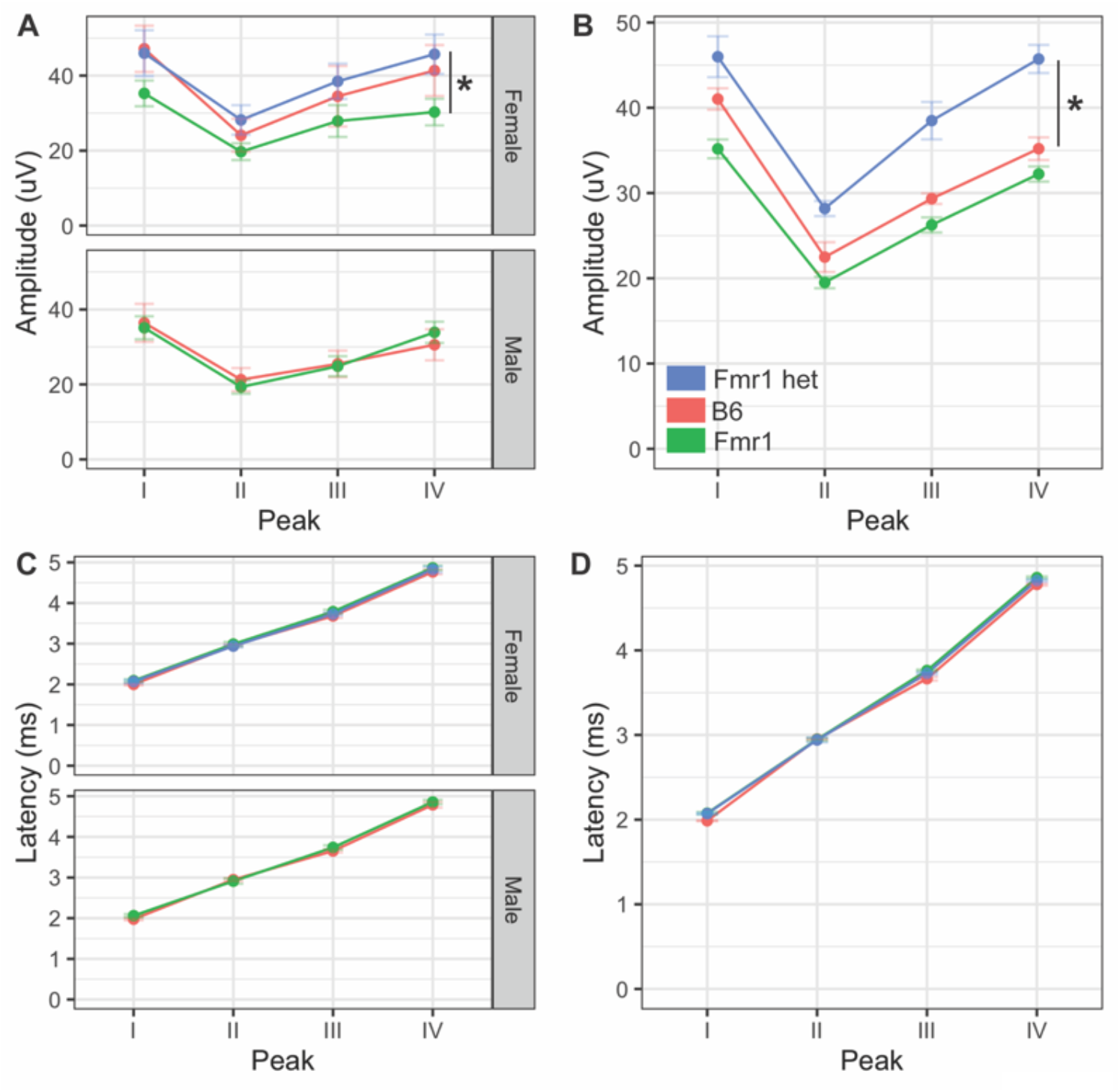
Monaural hearing in FXS mice. Monaural amplitudes and latencies for peaks I-IV of the ABR were recorded for Fmr1, Fmr1 het, and B6 animals. Peak IV amplitude was significantly lower in Fmr1 mice females compared to Fmr1 het females (A upper), There were no significant differences in amplitudes for males (A, lower). When combined, there was a significant difference in Fmr1 het animals compared to Fmr1 (B). There was no difference in latency of peaks I-IV between sexes (C) or genotypes (D). * = p < 0.05. Data represent 6 B6, 12 Fmr1, 9 Fmr1 het females and 8 B6, 14 Fmr1 males.

**Figure 5.**
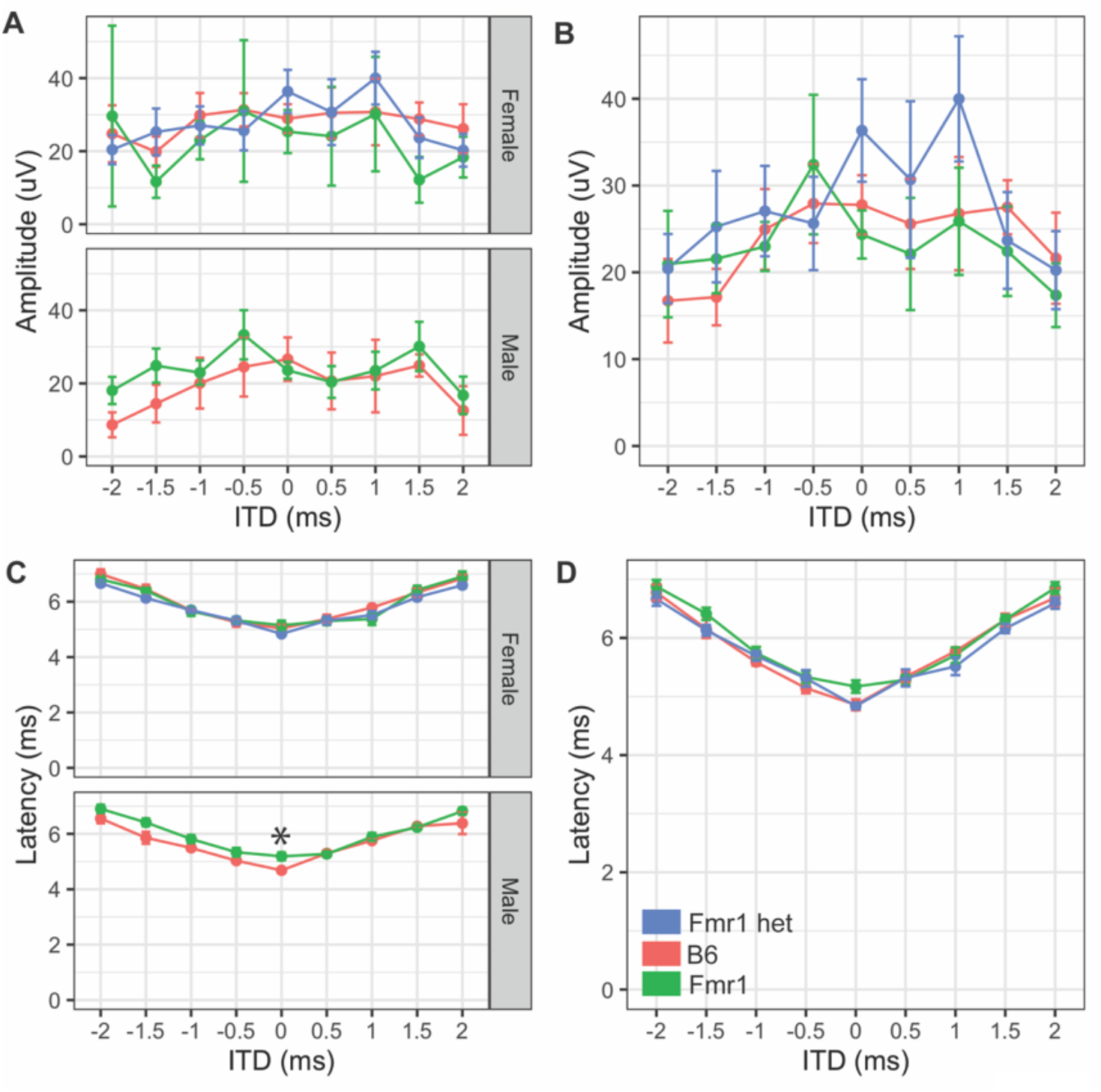
Binaural hearing in FXS mice. Binaural amplitudes and latencies for the BIC at ITDs between -2 to +2 ms in 0.5 ms steps were recorded for Fmr1 (green), Fmr1 het (blue), and B6 (red) animals. No differences in amplitude of the BIC with ITD for females (A upper), or males (A lower). When sexes were combined, there was no significant difference in amplitude of the BIC with ITD (B). Fmr1 males had significantly longer latency of the BIC at 0 ITD compared to B6 males (C, lower), while there was no difference in latency of female responses (C, upper). When sexes were combined, there was no difference in the BIC latency across ITDs between the genotypes (D). * = p < 0.05. Data represent 6 B6, 7 Fmr1, 9 Fmr1 het females and 6 B6, 9 Fmr1 males.

## 4 Results

We used both morphological and physiological features to examine hearing differences in a commonly used mouse model of FXS, C57BL/6J across genotypes and sexes. Hearing measurements included frequency hearing range, monaural hearing ability, and binaural processing using the ABR, while morphological features included pinna and head measurements.

### 4.1 Morphological features

People with FXS have altered craniofacial features, including large ears (Loesch et al., 1988). Consistent with our previous work (McCullagh et al., 2020a) we see no difference between B6, Fmr1, or Fmr1 het animals for pinna attributes (Figure 2C pinna width, 2D pinna length, 2B effective diameter). In addition, pinna characteristics were the same between the sexes (p = 0.175 pinna width, p = 0.96 pinna length, p = 0.267 effective diameter Figure 2B-D). When genotypes were compared within the same sex, there were no differences in weight, but sexes were significantly different independent of genotype (p = 0.0023) with females weighing significantly less than males. Similar to pinna morphology, there was no significant difference in either distance between pinna or distance from the nose to pinna between the genotypes or sexes (Figure 2E and F). These data suggest that mice do not share the same craniofacial changes, at least in the measurements described here, as people with FXS.

### 4.2 Hearing range

Our previous work showed that Fmr1 mice have increased thresholds for high frequency hearing compared to B6 at 16 kHz (McCullagh et al., 2020a). However, that work was limited by measuring only three frequencies (4, 8, 16 kHz) and seven mice of each genotype (combined sexes). Mice hear much higher frequencies than humans (Radziwon et al., 2009), therefore we wanted to measure whether this high frequency hearing loss exists across the frequencies in which mice hear in Fmr1 mutants and with a more in-depth sex specific analysis. Interestingly, there were no differences between genotypes across most of the frequencies tested except for an increase in hearing threshold at 64 kHz in Fmr1 mice compared to B6 (Figure 3B). There were no significant differences in hearing range between the sexes, but Fmr1 males did have significantly higher threshold at 64 kHz than female Fmr1 mice suggesting that the phenotype is mostly driven by males (p = 0.0317) and indeed both female Fmr1 and female Fmr1 het mice were not different than B6 females (Figure 3A). Best frequencies for both genotypes, as indicated by lower threshold, of mice were between 8 – 64 kHz consistent with specialized high frequency hearing.

### 4.3 Monaural hearing

Amplitude and latency of monaural ABRs correspond with neural activity across the ascending auditory pathway, with each wave representing different brain areas involved in auditory processing (Alvarado et al., 2012). Other studies have shown both latency and amplitude alterations in the FVB mouse strain of Fmr1 mutation (Kim et al., 2013; Rotschafer et al., 2015; El-Hassar et al., 2019). We measured ABR responses of Fmr1 mutants to monaural click stimuli compared to B6 mutant mice to determine if they have a similar ABR phenotype to the FVB strain. We saw no differences in overall click threshold for either genotype or sex (p = 0.102 genotype and p = 0.47 for sex). Amplitude of monaural responses was significantly lower for wave IV of the ABR in Fmr1 females compared to Fmr1 het females (Figure 4A upper). Indeed, Fmr1 het female amplitudes were closer to B6 than Fmr1 females, though Fmr1 females were not significantly different from B6. In contrast, Fmr1 male amplitudes for waves I-IV were not different from B6 (Figure 4A lower). When sexes were combined, Fmr1 het females had significantly higher amplitudes than B6, and were close to being significantly higher than Fmr1 mice (p = 0.0593). Consistent with sex driving the differences in genotype, peak amplitudes varied between the sexes. Female B6 mice had significantly higher amplitude peaks I and IV compared to B6 males (p = 0.0295 peak I and p = 0.0289 peak IV). In contrast, there were no sex differences between male and female Fmr1 mice suggesting a more male-like phenotype (independent of genotype) in homozygous Fmr1 females. There were no differences between the sexes or genotypes in latency of monaural peaks (Figure 4C and D).

### 4.4 Binaural hearing

While the monaural ABR provides information about binaural areas of the brainstem (potentially peaks III and IV), since they are elicited by either sound played directly to one ear (closed field) or equally to both ears (open field), little information can be gained about binaural integration of sound information. We used the BIC of the ABR to measure binaural processing ability of the brainstem as the BIC varies with ITDs played to both ears. We saw no differences in amplitude of the BIC at any ITD between the two genotypes (p = 0.809) or with sex (p = 0.6904, Figure 5A, B), though there was a significant difference between Fmr1 male and female mouse BIC amplitudes at 1.5 ms ITD. Latency of the BIC was significantly slower in male Fmr1 compared to B6 (Figure 5C, lower panel) only at 0 ITD, with no difference in genotype for female mice (Figure 5C, upper panel). When data were combined for sexes across genotype, there was no significant difference in latency of the BIC at any ITD (Figure 5D). There were differences in latency of the BIC between B6 (−1.5 ms) and Fmr1 (1 ms) males and females though there was no overall main effect of sex (p = 0.3367).

## 5 Discussion

This is the first study to characterize the ABR in the C57BL/6J Fmr1 mutant mouse, and in particular highlights morphological characteristics, hearing range, monaural ABRs, and binaural integration across sexes and in heterozygote females. Consistent with previous work, we see an increase in hearing threshold at high frequencies in Fmr1 mice, though this phenotype is male specific, and no change in morphology (pinna or facial characteristics)(McCullagh et al., 2020a). Female Fmr1 mice have reduced wave IV amplitudes of the monaural ABR, and wildtype females have increased wave I and IV amplitudes compared to B6 males, suggesting that female Fmr1 mice have a more male-like phenotype for monaural ABR amplitude. Lastly, we showed that male Fmr1 mice have increased latency of the BIC at 0 ITD, but not other ITDs or changes in amplitude of the BIC across ITD compared to B6 animals suggesting changes in timing of the processing of binaural information that does not change overall ITD following ability.

Pinnae size and shape are the first feature available to determine sound localization ability in animals with external ears (Butler, 1975; Musicant and Butler, 1984). Craniofacial alterations including prominent ears and elongated face are hallmark features of humans with FXS (Loesch et al., 1988; Heulens et al., 2013) and indeed may be a factor in auditory hypersensitivity that has been underexplored. Consistent with our previous work, we see no alterations in pinna or facial characteristics in the C57BL/6J mouse model of FXS (McCullagh et al., 2020a) using calipers as a measurement tool. Others have explored morphological skull differences in FXS mice using different tools such as CT/MRI (Ellegood et al., 2010) and micro-CT (Heulens et al., 2013) with mixed results. Heulens et al., 2013 showed alterations in skull and jaw characteristics that had not been characterized previously with a similar technique (Ellegood et al., 2010) though differences may be due to how features were measured. We also see no difference in weight of Fmr1 animals compared to wildtype which is in contrast to our previous work where we noted that Fmr1 animals weighed less than wildtype (McCullagh et al., 2020a) and others that showed an increase in male Fmr1 mouse weight compared to wildtype (Leboucher et al., 2019). Differences in weight may be due to inclusion of female animals (McCullagh et al., 2020a) and older animals (Leboucher et al., 2019). Overall changes in pinna morphology may still be an important factor in sound localization ability in Fmr1 animals and should be explored with more detailed techniques to determine if increased pinna measures in both humans and animal models may underly some aspect of auditory hypersensitivity symptomology.

Our previous results showed increased ABR measured hearing thresholds at high frequencies (16 kHz) in the C57BL/6J Fmr1 strain with data combined for the sexes (McCullagh et al., 2020a). In the current study, we do not see increased thresholds at 16 kHz but do see similar increased thresholds at 64 kHz in male Fmr1 mice specifically. These data are consistent with the increased thresholds across frequencies seen in adult male FVB Fmr1 mice (Rotschafer et al., 2015), though note that there was no change in threshold across frequencies in males of the same FVB strain at younger ages (Kim et al., 2013; El-Hassar et al., 2019). These data suggest that there may be age-related changes in high frequency hearing in adult male mice with FXS mutations across strains. This is particularly interesting since FVB mice are largely protected from early onset age-related hearing loss that the C57BL/6J background strain is particularly known for (Ison et al., 2007; Ho et al., 2014) suggesting that high frequency hearing deficits in males is a potentially conserved trait in FXS independent of background strain. Additional studies should examine hearing range across development and sexes in both strains to further show whether loss of high frequency hearing is a conserved feature in FXS.

Previous studies in the FVB Fmr1 mouse line show a robust wave I amplitude decrease in males across ages (Rotschafer et al., 2015; El-Hassar et al., 2019), though see (Kim et al., 2013). We do not see any change in wave I amplitude in the C57BL/6J Fmr1 line in adult animals of either sex. These conflicting results may be in part due to the earlier onset age-related hearing loss, which can be seen as decreases in early waves of the ABR, that occurs in the B6 background (Hunter and Willott, 1987). Changes in wave I amplitude specific to FXS may be masked by overall decreases in wave I amplitude across genotypes in this background. Interestingly, data in male FVB Fmr1 mice show no differences (adults, Kim et al., 2013; Rotschafer et al., 2015), or increased amplitudes in wave IV of the ABR (young, El-Hassar et al., 2019) whereas our data show decreased wave IV amplitude in Fmr1 females on the B6 background. These differences again may be due to differences in sexes and ages of animals tested. Lastly, our finding of no difference in latency of monaural waves is consistent with the majority of the work in FVB mice (Rotschafer et al., 2015; El-Hassar et al., 2019), though note that Kim et al., 2013 showed shorter latency for wave I. Our data further adds to the knowledge of ABR phenotypes that might be consistent across genotypes.

While ours is the first study to characterize the BIC in a FXS mutant mouse strain, our data are consistent with the BIC as it varies with ITD in mice (Benichoux et al., 2018). Namely, mice have a small range of ITD cues available due to their small head size and therefore the BIC amplitude decreases with increasing ITD between the ears, but this overall amplitude change is smaller than animals with more dominant ITD hearing ability (such as chinchilla or cats)(Benichoux et al., 2018). Additionally, consistent with previous work, the BIC latency gets longer with increasing ITD (Ferber et al., 2016; Laumen et al., 2016; Benichoux et al., 2018). Interestingly our work in FXS mice is consistent with increased latency of the BIC seen in a study in autistic people (ElMoazen et al., 2019), though they also see a decrease in amplitude of the BIC. Our findings that the BIC latency is only significant in males at 0 ITD potentially suggests that there is overall slowing of binaural processing in the brainstem, but that it is not dependent on ITD, which would be consistent with mice that do not rely as predominantly on ITD cues compared to other species.

The subject of sex differences in animal models is important for fully understanding the complexities of disorders such as autism spectrum disorder or FXS which seem to impact females differently than males (Werling and Geschwind, 2013; Nolan et al., 2017). In FXS, due to it being an X-linked disorder, there is a higher prevalence in males than females, which can undergo X-inactivation on the effected X chromosome (genetic mosaicism)(Kirchgessner et al., 1995). However, mice offer a unique opportunity to measure both heterozygote and homozygous females giving insight into potential sex differences related to loss of Fmr1 on one or both X chromosomes. Our data suggest that there are indeed differences in auditory phenotypes between heterozygous and homozygous females (wave IV amplitude) in addition to differences between males and females. These and future data comparing female Fmr1 subtypes may give insight into the role of X-inactivation in auditory brainstem processing phenotypes.

In conclusion, this study offers important insight into auditory phenotypes that may be shared or differ between background strains of FXS mice. Additionally, while subtle, we show sex-specific and full or heterozygote mutation-specific differences in auditory brainstem function for both monaural and binaural hearing in B6 mice. Further studies measuring auditory phenotypes for B6 mice in earlier ages across the sexes would be useful to further characterize potential similarities with the FVB Fmr1 strain. In addition, characterizing the BIC in the FVB strain would be useful to elucidate if latency phenotypes are consistent across backgrounds.

## 7 Author Contributions

All authors helped write and revise the manuscript. EAM developed the ideas and methods. EAM and AC collected the data for the manuscript. EAM performed the statistical analyses and created the figures for the manuscript.

## 9 Acknowledgements

We would like to thank members of the McCullagh lab on team mouse that assisted with ABRs including Ishani Ray and Sabiha Alam. Further we would like to acknowledge Shani Poleg and Daniel Tollin for helping us set up these experiments in Colorado and continue them in Oklahoma.

